# scTrends: automated classification and strength quantification of gene expression trends along pseudotime in single-cell RNA-seq

**DOI:** 10.64898/2026.04.21.719599

**Authors:** Jianbo Qing, Jiaying Hu, Xiao Wang, Junnan Wu

## Abstract

**Background:** Pseudotime inference has become a standard approach for reconstructing dynamic biological processes from single-cell transcriptomic data. However, after a pseudotemporal ordering has been established, systematically identifying and interpreting gene expression trends along pseudotime remains challenging. Existing approaches often rely on clustering-based heuristics or subjective parameter choices, which can compromise interpretability, reproducibility, and scalability in large datasets.

**Results:** We present *scTrends*, an automated and interpretable framework for gene-level trend classification and strength quantification along a given pseudotime trajectory. Importantly, *scTrends* does not perform pseudotime inference; instead, it operates downstream of established pseudotime methods to characterize expression dynamics once a temporal ordering is available. *scTrends* models pseudotime-binned gene expression profiles using generalized additive models and assigns genes to predefined temporal trend categories through a hierarchical, rule-based procedure combined with empirical significance testing, data-adaptive parameter selection, and quantitative assessment of trend strength. This enables simultaneous identification of the direction, shape, and magnitude of gene expression changes along pseudotime. We applied *scTrends* tothree distinct datasets: human PBMC, human brain, and mouse pancreas, using three different pseudotime inference methods (*CytoTRACE v2, Monocle3*, and *scVelo*, respectively). *scTrends* systematically characterized gene expression dynamics during T cell differentiation, oligodendrocyte precursor differentiation, and pancreatic endocrine cell maturation. The analysis revealed diverse monotonic, non-monotonic, and complex expression patterns, with varying strengths, consistent with known biological processes. Benchmarking analyses further demonstrate that *scTrends* is computationally efficient and scalable to large single-cell datasets, with modest memory requirements, making it suitable for diverse applications across a range of biological systems.

**Conclusions:** *scTrends* provides a systematic, automated, and resource-efficient solution for gene-level trend analysis in single-cell pseudotime studies, enabling reproducible characterization of dynamic expression patterns across diverse biological systems.

## 1. Background

The rapid development of single-cell RNA sequencing (scRNA seq) technologies has enabled a more detailed characterization of cellular and genetic changes during disease initiation and progression^1^. Dynamic alterations in cellular states and gene expression accompany the entire continuum of biological development, ageing, and disease. Dissecting cell differentiation trajectories and gene expression dynamics across diverse biological contexts has therefore become central to understanding life processes and identifying potential disease targets^2^.

Pseudotime inference methods, including *Monocle*^*3*^, *CytoTRACE*^*4*^, and RNA velocity-based approaches such as *velocyto*^*5*^, have become widely used for reconstructing continuous cellular trajectories from scRNA-seq data. These methods enable the ordering of cells along inferred developmental or differentiation axes and provide an essential foundation for studying dynamic biological processes. However, pseudotime inference alone does not address how gene expression dynamics should be systematically characterized and interpreted once a temporal ordering has been established.

Despite the availability of pseudotime estimates, accurately identifying, classifying, and quantifying gene expression trends along pseudotime remains challenging. Existing approaches typically rely on clustering-based strategies (for example, MUZZ, hierarchical clustering, or k-means) or curve-fitting frameworks such as tradeSeq to visualize pseudotime-dependent expression patterns^6^. While these methods are useful for smoothing expression trajectories or grouping genes with broadly similar temporal behaviors, they often require manual inspection to interpret specific dynamics and do not provide explicit, data-driven classifications of gene expression trends. Consequently, it can be challenging to objectively distinguish monotonic increases or decreases from transient peak-like, valley-like, or more complex multi-phase patterns, and genes with stable expression are rarely identified in a systematic manner. Moreover, many existing approaches lack formal statistical measures to assess whether observed trends are significantly distinct from noise, which can limit reproducibility and biological interpretability^7,8^.

To address these challenges, we developed *scTrends*, an R package designed for post-pseudotime gene expression trend analysis. *scTrends* operates downstream of established pseudotime inference methods and does not perform cell ordering or trajectory reconstruction. Instead, given a pseudotime axis, scTrends systematically identifies, classifies, and statistically evaluates gene expression dynamics using data-adaptive modeling and empirical significance testing. It categorizes genes into six biologically interpretable categories (Stable, Up, Down, UpDown, Down–Up, and Complex) while providing a quantitative trend strength score (0-1) that measures how typical a gene is of its assigned pattern and an omnibus permutation test for assessing overall temporal signal. By explicitly bridging pseudotime ordering with objective, reproducible gene-level dynamics, scTrends fills a critical methodological gap in single-cell trajectory analysis and facilitates more interpretable and biologically grounded downstream insights.

## 2. Implementation

### 2.1 *scTrends* overview

We developed *scTrends*, a comprehensive R package designed to automate the identification, classification, and statistical validation of gene expression trends along pseudotime trajectories in scRNA-seq data. Unlike existing tools that rely on predefined models or manual parameter tuning, *scTrends* adopts a data-adaptive framework that automatically optimizes classification criteria based on the characteristics of each dataset, thereby ensuring robustness across diverse biological systems and experimental platforms.

*scTrends* operates strictly as a downstream method and does not perform trajectory inference. Instead, it takes as input any user-provided continuous ordering of cells, represented as a numeric vector along a biological continuum. This ordering may be derived from linear pseudotime, RNA velocity-based time, diffusion pseudotime, or developmental potential estimated by methods such as *CytoTRACE v2*. This design ensures broad compatibility with existing trajectory inference frameworks while minimizing reliance on user-specified assumptions.

*scTrends* is fully implemented in R (version ≥ 4.1.0) and builds upon several core packages: 1) *mgcv* (v1.8-40 or later) for generalized additive model (GAM) fitting with automatic smoothing selection; 2) *parallel* and *pbapply* for efficient multicore computation and progress tracking; 3) base R statistical functions for permutation testing and multiple-testing correction.

The *scTrends* workflow consists of three main steps. First, data preprocessing aggregates gene expression into pseudotime bins to reduce noise and computational burden. Second, adaptive parameter selection employs quantile-based strategies and generalized cross-validation (GCV) minimization to optimize GAM smoothing and classification thresholds in a data-driven manner. Third, trend classification and statistical testing assign each gene to one of seven biologically interpretable categories (stable, borderline stable, up, down, up–down, down–up, and complex). Fourth, dual assessment is performed via trend strength scores (0-1, measuring pattern typicality) and omnibus permutation tests (evaluating statistical significance of temporal signals). This framework enables flexible filtering strategies based on either biological effect size or statistical confidence. Additionally, for datasets with branching structures, we recommend analyzing each lineage separately by subsetting cells corresponding to individual branches and applying scTrends independently. (**Figure 1**).

**Figure 1.**
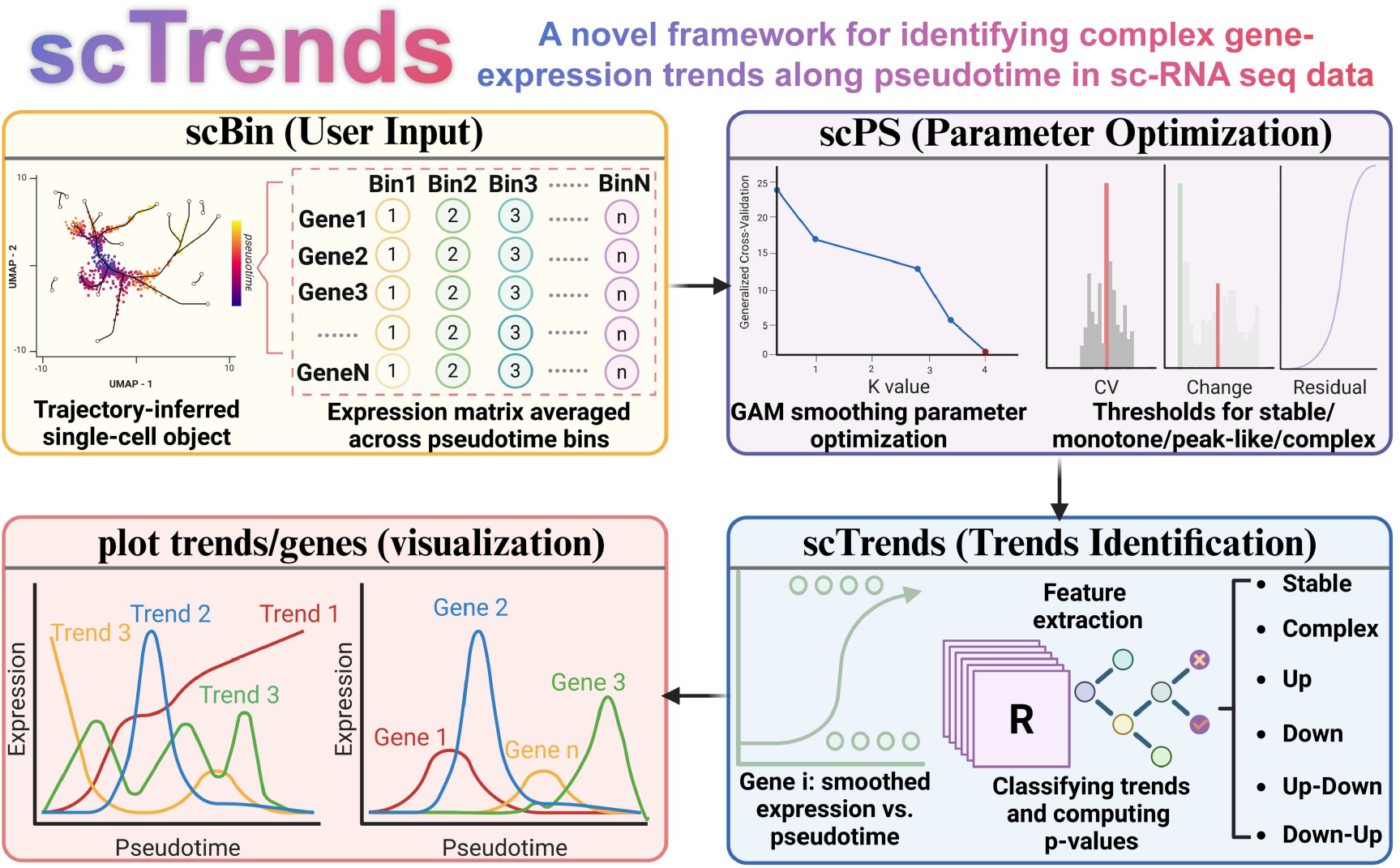
Comprehensive workflow of *scTrends*. The *scTrends* framework processes trajectory-inferred scRNA-seq data by first binning cells along pseudotime and averaging gene expression (scBin), then optimizing smoothing parameters and classification thresholds in a data-driven manner (scPS). It next fits expression curves to extract features, classify genes into distinct trend patterns, and assess statistical significance using permutation testing, with built-in functions for visualizing gene expression trajectories and representative trends along pseudotime. Abbreviations: scRNA-seq, single-cell RNA sequencing; GAM, generalized additive model; CV, coefficient of variation.

Comprehensive documentation, tutorials, and example datasets for *scTrends* are available at https://github.com/746443qjb/scTrends. *scTrends* supports single-cell data generated from major platforms, including 10x Genomics, Drop-seq, Smart-seq2,, and is compatible with widely used preprocessing workflows such as *Seurat* and *Scanpy*.

### 2.2 Data Preprocessing via Pseudotime Binning

scRNA-seq data are characterized by substantial technical noise arising from low capture efficiency, amplification bias, and dropout events^9^. When gene expression dynamics are analyzed along pseudotime trajectories, such noise can obscure genuine biological trends and substantially increase the computational burden of downstream modeling^8,10^. To address these challenges, *scTrends* adopts a pseudotime binning strategy that aggregates cells into discrete intervals, thereby reducing stochastic variation while preserving temporal resolution^11^. Cells are partitioned into a user-specified number of non-overlapping (or optionally overlapping) bins along the pseudotime axis, and gene expression values within each bin are averaged. The pseudotime coordinate for each bin is taken as its midpoint. This approach is computationally efficient, fully reproducible, and less sensitive to initialization or uneven cell density compared to clustering- or sliding-window-based smoothing (**Table 1**). As a practical guideline, we recommend 5-10 bins per major cell state and an average of at least 20 cells per bin to ensure stable mean estimation.

**Table 1.** Comparison of pseudotime smoothing strategies.

| Method | Strategy | Pros | Cons |
| --- | --- | --- | --- |
| No binning | Fit GAM on all cells | Maximum temporal resolution | High noise, slow computation |
| Moving average | Sliding window smoothing | Flexible resolution | Sensitive to uneven cell density |
| K-means binning | Cluster cells along pseudotime | Adaptive bin sizes | Sensitive to initialization |
| scTrends binning | Equal-width pseudotime intervals | Fast, reproducible | Fixed bin sizes |

To further balance noise reduction and temporal resolution, the “scBin” function optionally allows adjacent bins to share a specified proportion of cells, enabling smoother transitions between neighboring pseudotime intervals. In addition, genes whose expression values are zero in more than a user-defined fraction of bins can be filtered out prior to downstream analysis. These design choices jointly improve the robustness of trend estimation, reduce the influence of sparsity-driven artifacts, and enhance the statistical power of subsequent modeling steps.

As a practical guideline, we found that assigning approximately 5–10 bins per major cell state provides sufficient resolution to capture within-state variation and state-to-state transitions, while avoiding excessive fragmentation. In addition, to ensure stable estimation of bin-level means in sparse scRNA-seq data, we recommend maintaining an average of at least 20 cells per bin.

### 2.3 Adaptive parameter selection

Trend classification in *scTrends* depends on several decision thresholds (for example, criteria for stability, monotonicity, peak/valley patterns and complex dynamics). Because these thresholds are sensitive to dataset-specific properties such as sparsity, noise level and dynamic range, *scTrends* implements an automatic parameter selection module (“scPS” function) that derives data-adaptive thresholds from a representative subset of genes. The procedure comprises three steps: stratified gene sampling, optimization of the GAM basis dimension, and quantile-based threshold derivation from summary features. Hierarchical gene subsampling is recommended only when computational resources are limited or for preliminary testing.

#### 2.3.1 Optimization of GAM basis dimension

*scTrends* models binned expression trajectories using GAMs^12^. To select an appropriate basis dimension *k* for the spline smoother, “scPS” function evaluates candidate values

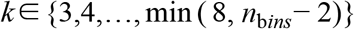

where *n*_b*ins*_ is the number of pseudotime bins. For each candidate *k*, a GAM is fitted to each sampled gene using pseudotime as the predictor, and GCV scores are computed^13^. The optimal basis dimension is chosen as

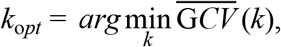

where 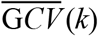 denotes the mean GCV score across sampled genes with valid fits. For datasets with limited numbers of bins (for example, when *n*_b*ins*_ −2 ≤ 3, scTrends defaults to *k*_o*pt*_ = 3).

#### 2.3.2 Feature extraction from fitted trajectories

Using *k*_o*pt*_, the “scPS” function fits a GAM to each sampled gene and extracts a set of summary features from the fitted trajectory 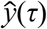 and residuals, including: (i) coefficient of variation (CV) and relative dynamic range of binned expression; (ii) lag-1 Spearman autocorrelation; (iii) peak/valley prominence relative to endpoints; (iv) normalized left/right slopes around extrema; (v) signed total change between trajectory endpoints; (vi) a monotonicity score based on the fraction of positive or negative first differences; (vii) the number of direction changes in first differences; and (viii) a residual-variance ratio quantifying unexplained variation. These features define empirical distributions that are subsequently used to derive classification thresholds.

#### 2.3.3 Quantile-based threshold derivation

Decision thresholds are derived from feature distributions using quantile rules that adapt to each dataset^14^. For stability detection, candidate stable genes are defined as those simultaneously within the bottom 20% for CV and relative change; thresholds are then set to the 95th percentile of these candidates for both CV and range. To reduce sensitivity to sparse candidates, *scTrends* requires at least 20 stable candidates; otherwise, conservative default thresholds are used. For autocorrelation, the stability threshold is defined using the lower tail of the high-autocorrelation subset (top 30%).

For peak/valley detection, prominence and slope thresholds are estimated from the lower tail of the high-prominence (above-median) and high-slope (above-median) subsets, respectively. For monotone trends, the signed total-change thresholds are derived from the 30th percentile of positive changes and the 70th percentile of negative changes, together with a monotonicity-consistency threshold set by the upper quantile of the monotonicity score distribution. For complex trends, the minimum number of direction changes is determined from the upper quantile of sign-change counts, and the residual-variance ratio threshold is derived from the upper quantile of residual-variance ratios. All thresholds are bounded within predefined ranges to prevent extreme values in small or noisy datasets.

### 2.4 Trend classification and statistical testing

*scTrends* assigns each gene to one of seven interpretable trend categories (Stable, Up, Down, Up–Down, Down–Up, and Complex) using a hierarchical decision procedure implemented in the “classify_single_gene” function. The method operates on binned expression profiles *g*_*m*_ defined over pseudotime bin centers *x* and combines rule-based pattern recognition with permutation-based significance testing.

In contrast to existing trajectory-based approaches that primarily rely on clustering or smooth curve fitting, *scTrends* directly focuses on trend-level characterization of gene expression dynamics along pseudotime^6,15,16^. Rather than requiring manual inspection of fitted curves or pre-selection of differentially expressed genes, *scTrends* performs automatic trend assignment for all genes based on explicit, interpretable criteria.

#### 2.4.1 Stable gene detection

A gene is classified as “Stable” if it satisfies a three-criterion stability test based on variability, dynamic range, and temporal coherence: (1) CV(*g*_*m*_) < *θ*_CV_ ; (2) 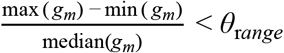; (3) *ρ*_*lag*1_(*g*_*m*_) > *θ*_*ac*_.

Where CV(*g*_*m*_) = sd(*g*_*m*_)/|mean(*g*_*m*_)|, and *ρ*_*lag*1_(*g*_*m*_) denotes the Spearman correlation between (*g*_1_,…,*g*_*n*−1_) and (*g*_2_,…,*g*_*n*_). In practice, the autocorrelation criterion is applied when n ≥ 3.

If permutation testing is enabled, *scTrends* computes an empirical *p*-value for stable genes using a residual-based statistic under a Gaussian noise null^17^. Specifically, a linear model is fitted to the observed profile,

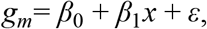

and the observed sum of squared residuals *S*_*obs*_ is compared with null residual sums *S*^*(b)*^ obtained by simulating 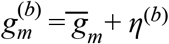, where 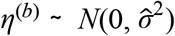. The empirical *P* value is computed as:

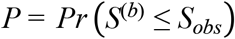

over *b* = 1,…, *B* simulations (default B = 1000, B is the total number of permutations).

#### 2.4.2 GAM fitting and global significance

For genes not classified as “Stable”, *scTrends* fits a GAM to capture smooth expression dynamics along pseudotime^13^:

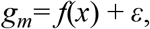

where *f*(.) is a spline smoother implemented via “mgcv::gam” function with basis dimension *k* = min(*k*_*opt*_, n - 2). The dataset-specific *k*_*opt*_ is obtained from “scPS” function.

Then, *scTrends* quantifies the strength of pseudotime dependence using the deviance-explained ratio:

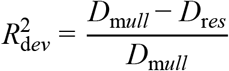

where *D*_m*ull*_ and *D*_r*es*_ are the null and residual deviances returned by the GAM fit. To assess statistical significance, an empirical *p* value is estimated using a permutation test^18^. Specifically, the binned expression values *g*_*m*_ are randomly permuted across pseudotime bins, thereby disrupting the temporal ordering, and the GAM is refitted to each permuted profile. Let 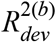 denote the deviance-explained ratio obtained from the *b*-th permutation (*b* = 1,…, B; default: B = 1000).

#### 2.4.3 Feature extraction

From the fitted trajectory *ĝ*(*x*) (the GAM-predicted values), *scTrends* computes a set of normalized features used for pattern classification:

1. **Prominence features**: peak and valley prominence relative to endpoint levels, normalized by the fitted range.
2. **Position features**: whether the global peak/valley lies away from the boundaries, defined by parameter “peak_position_margin”.
3. **Slope features**: left/right slopes around the peak (or valley), normalized by the fitted range.
4. **Monotonicity feature**s: signed net change between endpoints Δ = (*ĝ*(*x*_*n*_) − *ĝ*(*x*_1_) / r*ange*(*ĝ*) ; and 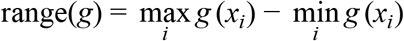 and first-difference consistency score max {*Pr* (Δ_*i*_ > 0), *Pr* (Δ_*i*_ < 0)}; and the number of sign changes in first differences.
5. **Complexity feature**: residual-variance ratio Var(*g*_*m*_−*g*) / Var(*g*_*m*_), and Var(·) denotes the sample variance over observed points.

#### 2.4.4 Hierarchical trend assignment

The hierarchical classification procedure follows a biologically motivated ordering. Stable expression is assessed first, as many genes exhibit minimal variation along a given trajectory. Transient patterns (Up-Down and Down-Up) are then prioritized, as they often reflect stage-specific regulatory events, such as transient activation of signaling or transcription factor networks. Monotonic trends are evaluated subsequently to capture gradual and sustained changes typical of differentiation processes. Finally, the remaining genes are assigned to “Complex” or “Borderline Stable” categories. This ordering not only improves computational efficiency but also reflects the increasing complexity of gene regulatory dynamics observed during development and disease processes. *scTrends* applies a prioritized rule set:

##### 2.4.4.1 Priority 1: Up–Down and Down–Up

A gene is classified as “Up–Down” if the fitted curve exhibits a central peak with sufficient prominence and opposing slopes, and all symbols *θ* denote user-specified thresholds controlling the stringency of pattern assignment. CV denotes the coefficient of variation of the fitted curve *g*_*m*_ :

1. peak prominence > *θ*_*prom*_ (the minimum normalized prominence required to classify a peak/valley as biologically meaningful).
2. peak index within the interior [*m, n* − *m*], *m* = [*n* × peak___position___margin].
3. left slope > *θ*_*slope*_ and right slope < −*θ*_*slope*_. *θ*_*slope*_ is minimum absolute slope threshold used to determine increasing or decreasing trends on either side of a peak or valley.
4. Analogously, “Down–Up” requires a central valley with prominence > *θ*_*slope*_ and slopes of opposite sign (left slope < *θ*_*slope*_ and right slope > −*θ*_*slope*_).

##### 2.4.4.2 Up and Down

If no peak/valley pattern is detected, *scTrends* evaluates monotone patterns. Let Δ denote the normalized signed net change and c the monotonicity consistency score. “Up” is assigned if:

1. Δ> *θ*_*up*_ (or relaxed threshold 0.8 *θ*_*up*_ when c > 1.15 *θ*_*cons*_). *θ*_*cons*_ is the minimum monotonicity consistency threshold, defined as the required proportion of first differences sharing the same sign.
2. c > *θ*_*cons*_
3. number of sign changes ≤ parameter “max_sign_changes_mono” (default: 1)
4. global slope sign > 0.

“Down” is defined analogously using *θ*_*down*_ < 0 and a negative global slope sign.

##### 2.4.4.3 Complex and Borderline Stable

If neither peak/valley nor monotone criteria are met, scTrends distinguishes “Complex” from “Borderline Stable”:

1. “Complex” if the direction-change count ≥ *θ*_*signchg*_ (maximum allowable number of sign changes in first differences for monotone classification.) or residual-variance ratio > *θ*_*var*_ (maximum residual-variance ratio threshold, defined as Var(gm − g) / Var(gm), beyond which trends are classified as complex.).
2. Otherwise, “Borderline Stable” is assigned to genes with weak net change and low variability:

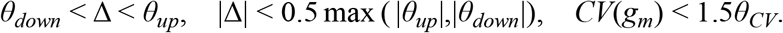

All remaining genes are labeled “Complex”.

Although the trend classification employs explicit rule-based thresholds for interpretability and reproducibility, it is complemented by two independent statistical/quantitative layers. The omnibus permutation test evaluates the global significance of temporal dependence (independent of category), while the trend strength score (0–1) quantifies how prototypical a gene is of its assigned pattern using type-specific saturating functions. Together, these provide both statistical grounding and practical confidence measures for downstream prioritization.

#### 2.4.5 Trend strength quantification

To quantify how typical a gene is for its assigned temporal trend pattern, we developed a trend strength scoring system based on saturating functions, yielding values between 0 and 1. Importantly, this score is independent of user-defined classification thresholds and is computed using trend-type–specific formulations.

For stable genes, strength reflects the degree of expression stability and is defined as:

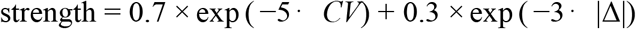

where CV denotes the coefficient of variation and Δ represents the normalized total expression change. Lower variability and smaller overall changes result in higher strength scores.

For monotonic genes (Up or Down), strength integrates both the magnitude of change and the consistency of monotonic behavior:

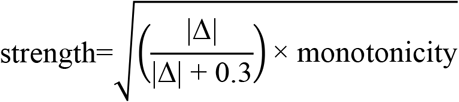

where |Δ| is the normalized net expression change and 0.3 serves as a half-saturation constant, analogous to Michaelis–Menten kinetics. *monotonicity* (range 0–1) quantifies the directional consistency of expression changes. A geometric mean is used to ensure that both components must be high to achieve a high strength score.

For peak–valley patterns (Up–Down or Down–Up), strength is determined by the prominence of the peak or valley:

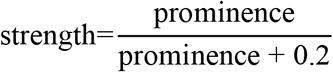

where *prominence* denotes the normalized peak/valley prominence and 0.2 is a half-saturation constant.

For complex patterns, strength captures oscillatory complexity and is defined as:

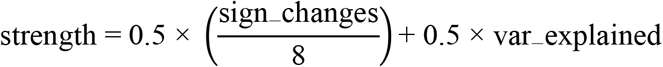

where *sign_changes* is the number of sign changes in the first derivative of the fitted curve, and *var_explained* represents the proportion of variance explained by the GAM.

This score serves as a continuous confidence measure for the assigned category and is independent of the decision thresholds, allowing users to filter or rank genes even when p-values are marginal due to data sparsity.

#### 2.4.6 Statistical significance testing

To assess whether a gene exhibits a statistically significant temporal signal, we employed an omnibus permutation test^19^. The null hypothesis assumes no temporal structure along pseudotime, whereas the alternative hypothesis posits the presence of any temporal pattern, without specifying a particular trend type.

For each gene, 1,000 permutations were performed by randomly shuffling the correspondence between expression values and pseudotime bins, thereby generating an empirical null distribution of the test statistic. *P* values were computed based on the rank of the observed statistic within this null distribution and could be adjusted for multiple testing using the Benjamini–Hochberg procedure (*FDR* < 0.05). Importantly, this *P* value tests whether a gene exhibits any temporal signal, rather than whether it belongs to a specific trend category. Trend classification itself is performed using a rule-based feature set.

The omnibus *P* value tests for the presence of temporal structure but does not directly reflect the typicality or biological cleanliness of a specific trend shape. Therefore, we recommend interpreting results jointly with the independent trend strength score, which quantifies how well a gene exemplifies its assigned pattern (e.g., high monotonicity consistency for Up/Down genes, or high prominence for transient patterns). This dual metric approach helps prioritize biologically meaningful genes even when p-values vary due to data sparsity after binning.

### 2.5 Visualization of Trends and Genes

To facilitate interpretation and downstream reporting, *scTrends* provides two lightweight visualization functions, “plot_trends” and “plot_genes”, both implemented using *ggplot2*. Because the returned objects are standard ggplot objects, users can freely customize aesthetics to match publication or presentation styles.

### 2.6 Data testing

To evaluate the performance of *scTrends*, we utilized the publicly available human PBMC (pbmcMultiome) dataset from 10x Genomics, along with datasets from the human brain and mouse pancreas. Pseudotime trajectories for the PBMC dataset were inferred using *CytoTRACE v2* (version 1.0.0), while *Monocle3* (version 1.3.4) was used for the human brain dataset and *scVelo* (version 0.3.4) for the mouse pancreas dataset. The resulting pseudotime values were then used for testing the scTrends package.

To benchmark *scTrends* classification against conventional unsupervised methods, we performed hierarchical clustering with Ward.D2 linkage and k-means clustering on the same binned pseudotime expression matrix for each dataset. Agreement between the scTrends assignments and clustering results was quantified using the Adjusted Rand Index (ARI) and Normalized Mutual Information (NMI), both implemented via the mclust and igraph packages in R.

All analyses were performed in R (version 4.3.1) and Python (version 3.10) on a workstation equipped with an Intel Core i7-13700KF processor.

## 3. Results

### 3.1 Pseudotime analysis and binning

To evaluate the compatibility and applicability of *scTrends* across different pseudotime inference methods and diverse datasets, we analyzed three distinct tissue datasets: the 10x Genomics PBMC reference dataset (pbmc3k), a human brain dataset, and a pancreas dataset. For pseudotime estimation, we applied *CytoTRACE v2* to the PBMC dataset, *Monocle3* to the brain dataset, and RNA velocity (*scVelo*) to the pancreas dataset. This multi-dataset, multi-method approach allows us to systematically assess scTrends’ performance in classifying gene expression trends across different biological systems and upstream pseudotime frameworks.

First, the 10X Genomics PBMC reference dataset contains pre-annotated peripheral blood mononuclear cell types, including T cells, B cells, and monocytes. To evaluate scTrends, we extracted CD4 naïve T cells and CD4 central memory T cells (CD4 TCM) to investigate gene expression dynamics during CD4 T-cell differentiation. A total of 2,568 CD4 T cells were retained for downstream analysis. Principal component analysis (PCA) was performed on the extracted cells, followed by visualization using Uniform Manifold Approximation and Projection (UMAP; **Figure 2A**). Differentiation potential was estimated using *CytoTRACE v2*, chosen for its objective, data-driven inference without requiring manual specification of a root cell. As expected, CD4 naïve T cells exhibited higher differentiation potential and were positioned at earlier pseudotime stages, whereas CD4 TCM cells occupied later stages (**Figure 2B and Table S1**). The trajectory was partitioned into 20 pseudotime bins using the “scBin” function, resulting in approximately 125 cells per bin. Genes with zero expression in any bin were excluded, leaving 9,702 genes for subsequent trend analysis. Expression patterns of the top 20 genes are shown as a heatmap in **Figure 2C**.

**Figure 2.**
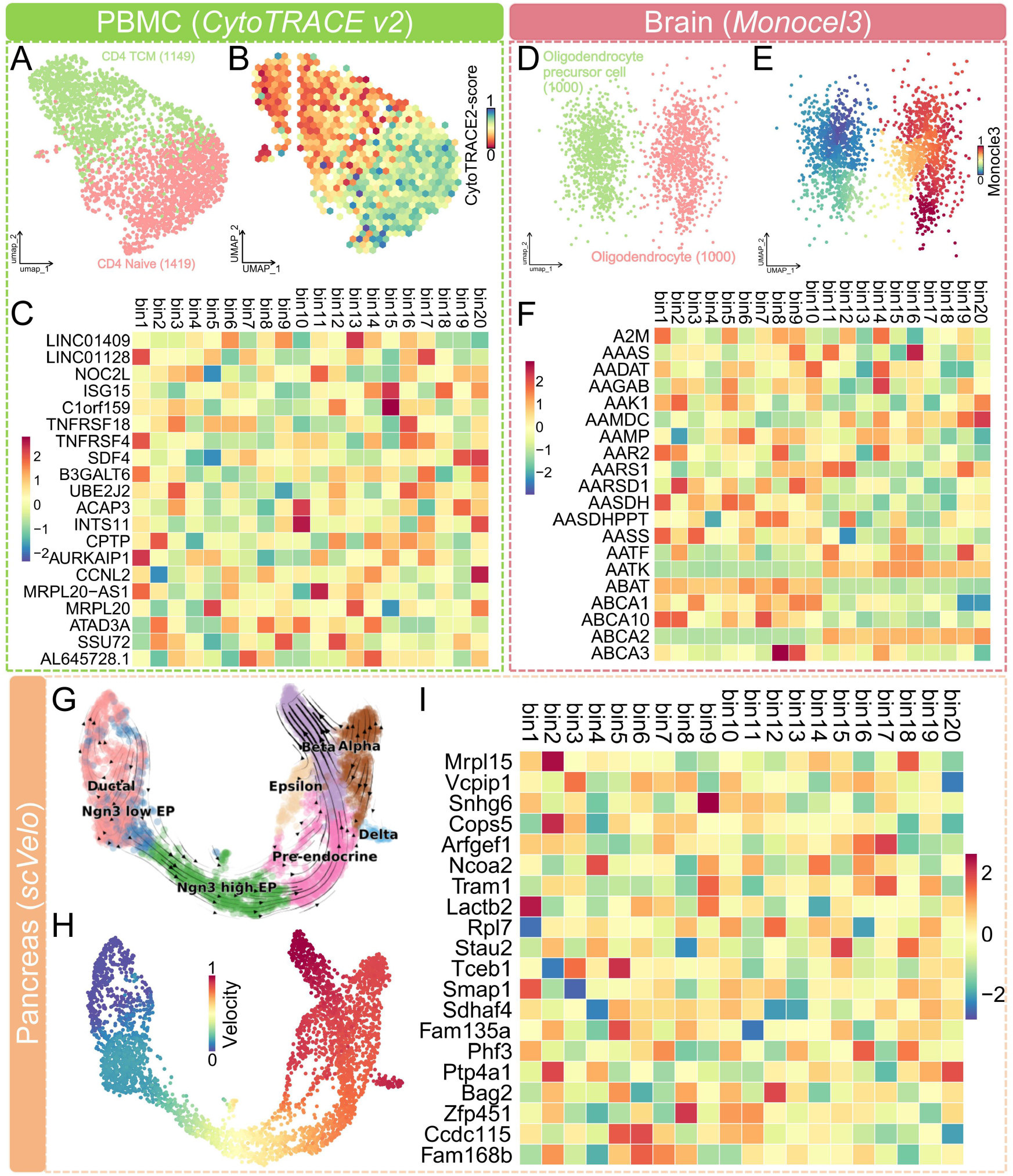
Pseudotime analysis and binning of three datasets. (A) UMAP visualization of CD4 naïve T and TCM cells. (B) Feature plot showing the differentiation potential (pseudotime ordering) of CD4 T cells inferred by *CytoTRACE v2*, where warmer colors indicate higher differentiation status and later pseudotime, while cooler colors indicate earlier pseudotime. (C) Heatmap of the top 20 expressed genes across the 20 pseudotime bins of PBMC dataset. (D) UMAP visualization of oligodendrocyte precursor cells and oligodendrocytes. (E) Feature plot showing the pseudotime ordering of oligodendrocytes inferred by *Monocle3*, where warmer colors indicate earlier pseudotime, while cooler colors indicate later pseudotime. (F) Heatmap of the top 20 expressed genes across the 20 pseudotime bins of brain dataset. (G) UMAP visualization of pancreatic cells. (H) Feature plot showing the RNA velocity of Pancreatic cells inferred by *scVelo*, where warmer colors indicate earlier pseudotime, while cooler colors indicate later pseudotime. (I) Heatmap of the top 20 expressed genes across the 20 pseudotime bins of pancreas dataset.

We next analyzed a dataset comprising 1,000 oligodendrocyte precursor cells and 1,000 oligodendrocytes (**Figure 2D**). Pseudotime was estimated using *Monocle3* **(Figure 2E and Table S2)**, and cells were partitioned into 20 pseudotime bins, yielding 8,158 genes for downstream analysis. The expression patterns of the top 20 genes are shown in **Figure 2F**.

Finally, we examined a mouse pancreas dataset, containing ductal cells, endocrine progenitors, Pre-endocrine, Beta, Alpha, Delta, and Epsilon cells (**Figrue 2G**). RNA velocity analysis (*scVelo*) was used to estimate pseudotime, which allows the modeling of branching differentiation trajectories (**Figure 2H and Table S3**). To focus on a single differentiation path, we selected Pre-endocrine (592 cells) to Beta (591 cells) cells. Cells were partitioned into 20 pseudotime bins, and 7,737 genes were retained for trend analysis. The top 20 genes are displayed in **Figure 2I**.

### 3.2 Automatic, dataset-specific calibration of *scTrends* parameters

Using the binned pseudotime expression matrices generated from each dataset, we applied the “scPS” function to obtain a dataset-specific set of parameters for downstream *scTrends* analysis. Notably, this step is optional: users can either manually specify all parameters or allow “scPS” to automatically compute dataset-adaptive values. In our analyses, all genes were included for parameter selection across the three datasets (**Figure S1-S3**).

To further assess the robustness of the dataset-adaptive parameter selection, we performed random subsampling of cells in each dataset (PBMC, oligodendrocyte precursor cells, and pancreas). Cells were sampled at 20%, 40%, 60%, and 80% of the original dataset size, with 10 independent runs per sampling proportion. For each of the eight key thresholds, we calculated the mean SD across runs and datasets.

As shown in **Figure 3**, the thresholds estimated by “scPS” exhibited generally low variability, which decreased with increasing sampling proportion. Parameters related to stable gene detection, such as range, autocorrelation, and consistency thresholds, showed minimal variation (mean SD < 0.015) and were largely insensitive to sampling proportion, indicating high intrinsic stability of these quantile-based rules. In contrast, total change thresholds for Up and Down trends displayed relatively higher variability at low sampling proportions (mean SD ≈ 0.035 at 20%) but decreased substantially with more cells included (mean SD < 0.012 at 80%). Remaining parameters (CV threshold, minimum prominence, and minimum slope) exhibited moderate variability that also decreased with increasing cell numbers.

**Figure 3.**
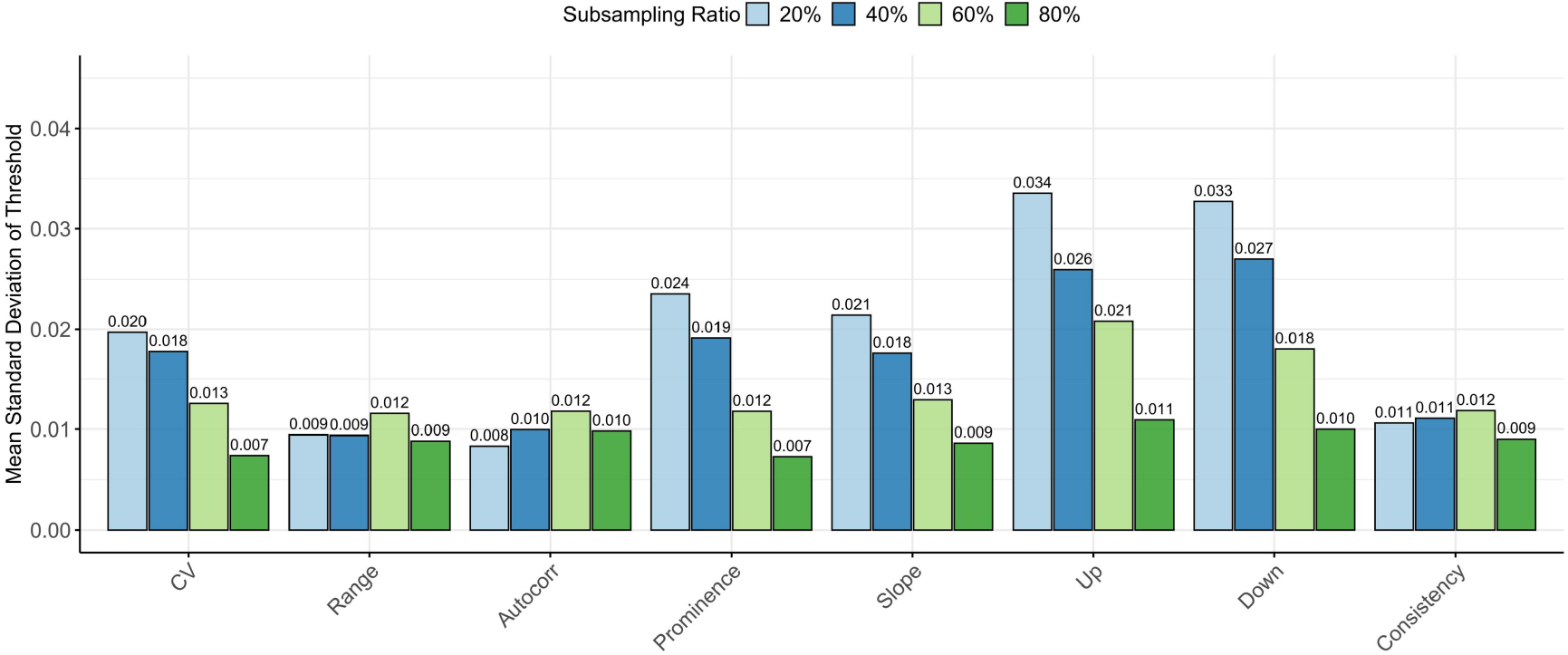
Robustness of scPS threshold selection under cell subsampling. We randomly subsampled cells at 20%, 40%, 60%, and 80% of the original size (10 independent runs per ratio) across three datasets (human PBMC, human brain, and mouse pancreas). For each parameter, the mean SD of the derived threshold across runs and datasets is shown. Parameters such as Consistency, Autocorr, and Range showed consistently low variability regardless of subsampling ratio, while Up and Down thresholds exhibited higher variability at low subsampling ratios that decreased markedly as more cells were included. Overall, variability remained modest even at 20% subsampling, supporting the robustness of the quantile-based, data-adaptive parameter optimization in “scPS”.

Overall, even at the most stringent 20% sampling level, threshold variability remained within biologically and statistically acceptable ranges. Nevertheless, hierarchical gene subsampling is recommended only when computational resources are limited or for preliminary testing, as the deterministic use of all genes provides maximal stability and reproducibility.

### 3.3 Classification of gene expression dynamics across datasets

Using scTrends, the 9,702 genes retained from the PBMC dataset were classified into five categories based on their expression dynamics along pseudotime: “Complex” (1,042 genes), “Up” (4,062 genes), “Down” (2,218 genes), “Up-Down” (1,348 genes), and “Down-Up” (1,032 genes), with no genes identified as stably expressed (**Table S4**). A similar predominance of dynamic trends was observed in the additional datasets. In the human oligodendrocyte lineage dataset, two genes were classified as “Stable”, whereas 2,444 were assigned to “Complex”, 1,886 to “Down”, 1,166 to “Down-Up”, 1,897 to “Up”, and 763 to “Up-Down” (**Table S5**). In the pancreas dataset, 814 genes were classified as “Complex”, 2,594 as “Down”, 870 as “Down-Up”, 2,360 as “Up”, and 1,099 as “Up-Down” (**Table S6**). Considering the representativeness of T-cell differentiation, we next conducted a detailed exploration of gene expression dynamics during this process.

**Figure 4A** shows the fitted expression trajectories of all 9,702 genes across 20 pseudotime bins, revealing highly dynamic and diverse gene expression changes during the differentiation of CD4 naïve T cells toward the TCM state. In addition, we visualized the fitted expression curves of the top 30 most significant genes (ranked by *P* value) from each of the “Up”, “Down”, “Up-Down”, “Down-Up” and “Complex” categories (**Figure 4B-F**). Overall, these genes displayed expression patterns that were highly consistent with their respective scTrends-defined trends, indicating that scTrends effectively captures the dynamic transcriptional changes occurring during T cell differentiation.

**Figure 4.**
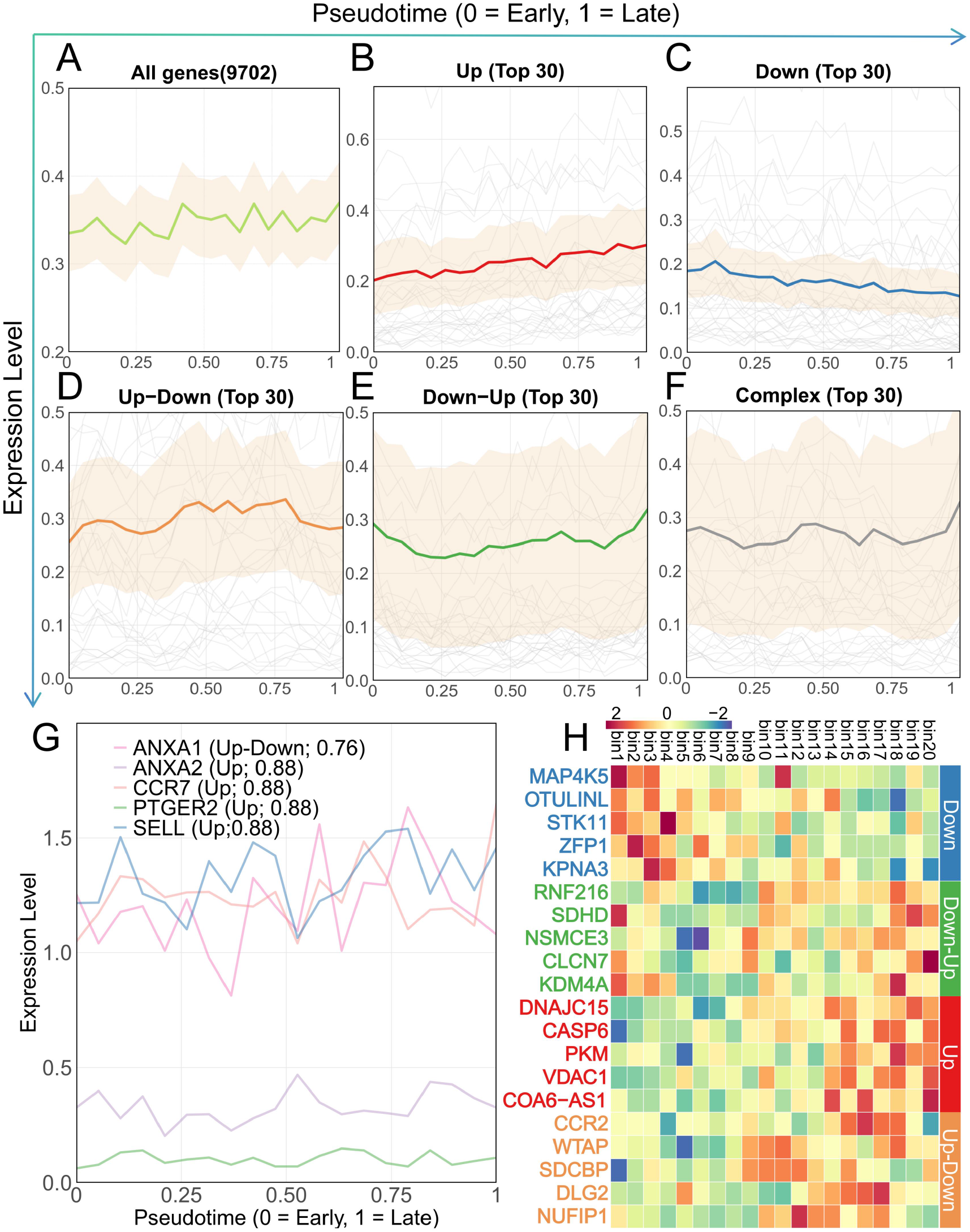
Identification of gene expression trends during CD4 naïve to TCM differentiation. (A) Fitted expression trajectories for all 9,702 genes after binning CD4 T cells along pseudotime. (B–F) Fitted expression curves of the top 30 most significant genes for each trend category: “Up”, “Down”, “Up-Down”, “Down-Up”, and “Complex”. Shaded regions represent approximate 95% confidence intervals around the fitted curves, calculated as ± 1.96 standard errors. (G) Pseudotime expression profiles of five representative TCM marker genes, illustrating their dynamic regulation during the transition from naïve CD4 T cells to the TCM state. (H) Heatmaps showing the expression and the trend strength of the top five most significant genes from the “Up”, “Down”, “Up-Down”, “Down-Up” categories across 20 pseudotime bins.

Previous work by Zhang et al. identified several characteristic markers distinguishing peripheral blood TCM cells from CD4 naïve T cells, including *CCR7, ANXA1, ANXA2, PTGER2*, and *SELL*^20^, and all of these genes show a trend strength greater than 0.7. Along the pseudotime trajectory constructed in this study, *CCR7, ANXA2, PTGER2*, and *SELL* were all classified as exhibiting monotonically increasing (Up) expression patterns, whereas *ANXA1* was identified as following an “Up-Down” pattern, characterized by an initial increase followed by a subsequent decrease in expression (**Figure 4G**). Consistent with their established roles, *CCR7* and *SELL*, which are key molecules involved in lymphoid homing of central memory T cells^21,22^, showed gradual upregulation during differentiation, in line with the acquisition of the TCM phenotype. *PTGER2* and *ANXA2*, which have been implicated in T cell migration, inflammatory regulation, and memory T cell function, also displayed increasing expression trends that agree with previous reports^23-25^.

Notably, *ANXA1* is known to be dynamically regulated during T cell activation and immune responses^26^, where it participates in TCR signaling and modulates proliferation and differentiation programs rather than being constitutively expressed throughout T cell states^27^. This supports the possibility that an initial increase followed by a decrease in *ANXA1* expression along pseudotime may reflect its stage-specific regulatory role early in differentiation, preceding the establishment of a mature TCM phenotype^28^.

To further evaluate the reliability of *scTrend* in identifying gene expression dynamics along pseudotime, we examined representative genes from each trend category. As shown in **Figure 4H**, we visualized the expression of the five most significant genes (ranked by *P* value) from each of the “Up”, “Down”, “Up-Down”, and “Down-Up” categories across the 20 defined pseudotime bins.

Genes classified as “Down”, such as *STK11* (*LKB1*) and *MAP4K5*, showed progressively decreasing expression along pseudotime. Both genes have been linked to T cell homeostasis and signaling, and their downregulation is consistent with a shift away from the naïve state during differentiation toward TCM^29,30^. The “Down-Up” category included genes such as *KDM4A* and *RNF216*, which are associated with chromatin regulation and protein ubiquitination^31,32^, respectively, and exhibited an initial decrease followed by re-upregulation along pseudotime. Genes classified as “Up”, including *PKM* and *VDAC1*, showed progressively increasing expression and are related to cellular metabolism and mitochondrial function, consistent with gradual metabolic remodeling during differentiation^33,34^. In contrast, “Up-Down” genes such as *CCR2* and *WTAP* displayed transient upregulation at intermediate pseudotime stages. *CCR2* has been linked to T cell migration^35^, whereas *WTAP* is involved in post-transcriptional regulation through m6A RNA methylation^36^.

Notably, the “Complex” category (1,042 genes) encompasses genes whose expression dynamics cannot be captured by simple monotonic or single-peak patterns, often exhibiting multiple directional changes or high residual variance, suggestive of multi-phase or oscillatory regulatory events. Representative genes in this category, such as *NR4A2, ATG5*, and *TNFSF10* have clear associations with immune regulation and T cell differentiation. *NR4A2*, a nuclear receptor transcription factor, can directly induce Foxp3 and regulate the Treg differentiation program in CD4+ T cells, suggesting its key role in the determination of regulatory T cell fate^37^. ATG5, as a core autophagy factor, has experimental evidence supporting its role in T cell survival and functional maintenance, revealing that autophagy may be involved in immune regulatory dynamics^38^. Additionally, TNFSF10 (also known as TRAIL), a member of the TNF family, is widely discussed in immune regulation and cell fate signaling^39^. These genes exhibit diverse temporal expression profiles, suggesting their involvement in complex multi-node regulatory programs during T cell differentiation

Additionally, for the human cortical oligodendrocyte dataset, we successfully identified genes such as *NEO1*, whose expression increases during the maturation of oligodendrocytes^40^. Similarly, in the pancreas dataset, the gene *Pdx1*, which is involved in the differentiation of Pre-endocrine cells into Beta cells, showed a progressive increase in expression during the later stages of differentiation^41^. These results collectively demonstrate the ability of *scTrends* to accurately capture gene expression dynamics across diverse differentiation trajectories, highlighting its robustness and adaptability in analyzing complex biological processes.

### 3.4 Benchmarking scTrends against conventional clustering methods

To evaluate the relationship between the trend categories derived from *scTrends* and traditional unsupervised gene grouping methods, we compared our classification results with hierarchical clustering (Ward.D2 linkage) and k-means clustering applied to the same binned pseudotime expression matrices from the three datasets.

Across all datasets, the numerical consistency between scTrends and clustering methods was low, with ARI values ranging from -0.0098 to 0.0065 and NMI values near 0 (**Table 2**). This low consistency was expected and highlights a key strength of scTrends rather than a limitation. Conventional clustering methods group genes based on overall expression pattern similarity (Euclidean distance), often resulting in the mixing of monotonic “Up” and “Down” genes due to their shape symmetry, disrupting transient “Up–Down” or “Down–Up” patterns, and scattering genes with multi-phase dynamics classified as “Complex.” In contrast, scTrends utilizes biologically-driven, rule-based criteria (derived from features such as peak prominence, slope direction, and residual variance from GAM fitting) to assign distinct and interpretable trend categories, accompanied by trend strength scores (0–1) and permutation-based statistical significance. Thus, *scTrends* serves as a valuable complement to conventional clustering, offering more actionable biological insights without the need for extensive manual post-annotation of clusters.

**Table 2.** Agreement between *scTrends* trend classification and conventional clustering methods.

| Dataset | Method | ARI | NMI |
| --- | --- | --- | --- |
| Human PBMC | Hierarchical (Ward.D2) | -0.0098 | 0.0104 |
| Human PBMC | k-means (k=5) | -0.0084 | 0.0102 |
| Human brain | Hierarchical (Ward.D2) | -0.0012 | 0.051 |
| Human brain | k-means (k=6) | 0.0065 | 0.0435 |
| Mouse pancreas | Hierarchical (Ward.D2) | 0.001 | 0.002 |
| Mouse pancreas | k-means (k=5) | 0.001 | 0.001 |

### 3.5 Computational time and memory usage

*scTrends* relies exclusively on CPU-based computation. All benchmarks were performed on a workstation equipped with an Intel Core i7-13700KF processor (16 cores) using parallel execution. We systematically evaluated the computational time and memory consumption of *scTrends* under different parameter settings, with three independent runs conducted for each configuration. All experiments were performed using a GAM with *k* = 8.

When *P*-value computation was disabled, *scTrends* showed high computational efficiency. Under a setting of 20 pseudotime bins and 10,000 genes, the classification step completed in approximately 1 minute. Varying the number of bins or genes had only a marginal impact on runtime in this mode, and therefore no further benchmarking was performed. In contrast, when *P*-values were computed using permutation testing (default number of permutations = 1,000), computational time and memory usage increased substantially. Under the baseline setting of 20 bins and 10,000 genes, *scTrends* required an average of 57 minutes and approximately 8 GB of random access memory (RAM).

Notably, both runtime and RAM usage increased approximately linearly with respect to the number of pseudotime bins and the number of genes (**Figure 5A-5D**). When the number of genes was fixed at 10,000, increasing the number of bins by 10 resulted in an average increase of approximately 50 minutes in runtime and 4 GB in memory usage. Conversely, when the number of bins was fixed at 20, increasing the number of genes by 10,000 led to an average increase of approximately 6 minutes in runtime and 0.8 GB in memory consumption.

**Figure 5.**
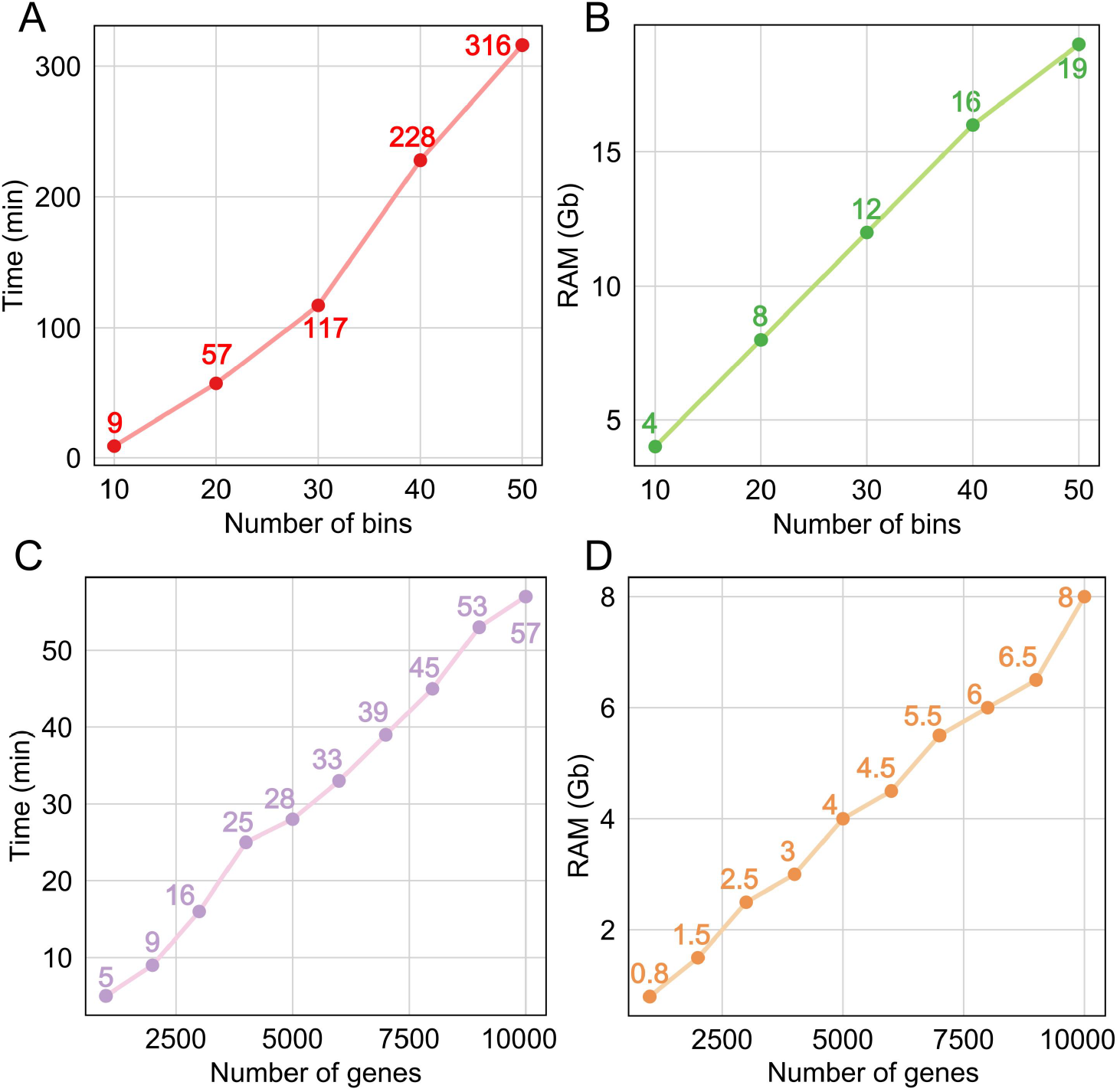
Computational performance of *scTrends* under varying bin and gene numbers. Runtime (A) and RAM (B) consumption of scTrends as a function of the number of bins with the gene number fixed at 10,000. Runtime (C) and RAM (D) consumption of *scTrends* as a function of the number of genes with the number of bins fixed at 20.

Under a configuration of 50 pseudotime bins and 10,000 genes, *scTrends* required an average runtime of 316 minutes with a peak memory usage of approximately 19 GB. This level of computational demand is readily manageable on both standard personal workstations and typical server environments. These results indicate that *scTrends* can be efficiently executed under typical parameter settings on commonly available hardware.

However, due to the rapid increase in model complexity and node number associated with higher bin counts, excessive bin partitioning substantially elevates computational cost and reduces practical efficiency. Based on these observations, we recommend using no more than 50 pseudotime bins in practical applications to achieve a balance between resolution and computational efficiency.

## 4. Discussion

In this study, we introduce *scTrends*, a computational framework specifically designed to automatically identify, classify, and statistically validate gene expression trends along established pseudotime trajectories in scRNA-seq data. Importantly, *scTrends* operates downstream of pseudotime analyses and does not perform trajectory inference itself, addressing key challenges in the systematic characterization of dynamic gene expression programs. While existing pseudotime inference methods provide effective tools for ordering cells along a pseudotemporal axis, they do not systematically classify gene expression patterns or quantify trend strength. *scTrends* bridges this gap by offering a rigorous framework to categorize gene expression dynamics into biologically interpretable patterns and to quantify their robustness. Furthermore, its design is compatible with velocity- or dynamically informed pseudotime, allowing integration with upstream dynamic modeling approaches while maintaining interpretability and reproducibility.

Traditional approaches to pseudotime-based gene expression analysis often rely on clustering or curve-fitting methods, such as GAMs. While these methods can generate smoothed trajectories or group genes based on overall expression patterns, they fail to provide clear and reproducible classifications of dynamic gene expression trends. Moreover, the interpretation of gene behavior along pseudotime is typically subjective, relying on manual inspection to identify trends like monotonic increases or decreases, transient upregulation, or oscillatory behavior. This subjectivity is especially problematic for identifying subtle or low-amplitude trends and genes exhibiting more complex or non-monotonic dynamics, which can be obscured by noise or sparsity in single-cell data.

*scTrends* addresses these challenges through a fully automated, rule-based framework. It first aggregates expression into pseudotime bins to reduce technical noise, followed by dataset-specific parameter optimization using generalized cross-validation and quantile rules (scPS). This is followed by GAM fitting, feature extraction, hierarchical trend classification (Stable, Up, Down, Up–Down, Down–Up, Complex), trend strength quantification, and permutation-based significance testing. Importantly, all parameters are fully customizable, while also incorporating data-driven parameter optimization to automatically determine key model parameters from the input data. This design ensures robust default settings while providing flexibility for advanced users, significantly improving reproducibility and minimizing reliance on subjective judgment.

We tested three datasets, each utilizing a different pseudotime inference method. This approach ensured compatibility across various methods and allowed us to effectively address cases involving branching trajectories. Furthermore, our results maintain strong interpretability, aligning with established biological processes and offering valuable insights into the dynamics of gene expression. For example, we used the differentiation process from CD4 naïve T cells to TCM as a representative biological process. *scTrends* accurately identified well-established genes exhibiting increasing expression during this process, such as *CCR7*, while also revealing additional genes with more complex dynamics, including *STK11* and *CCR2*, which may play previously underappreciated roles during T cell differentiation. Notably, the expression trajectories of these genes display substantial fluctuations along pseudotime, making reliable trend identification through manual inspection challenging. *scTrends* effectively disentangled these complex patterns, demonstrating its ability to recover biologically meaningful trends that would otherwise be difficult to discern.

In terms of computational efficiency, *scTrends* is lightweight and can be applied to large datasets with modest computational resources. The tool runs efficiently on standard personal workstations, demonstrating its scalability for larger single-cell transcriptomic datasets. By incorporating automated parameter optimization and trend strength quantification, *scTrends* offers an end-to-end solution for analyzing gene expression trends along pseudotime trajectories without requiring manual intervention or assumptions about trend shapes. This robust and flexible design positions *scTrends* as a powerful tool for post-pseudotime gene expression trend analysis, complementing existing pseudotime inference methods and enabling comprehensive, reproducible analyses of complex transcriptional dynamics in single-cell studies.

scTrends has a broad impact on single-cell trajectory analysis. First, it transforms pseudotime from a mere ordering tool into a framework for the quantitative analysis of dynamic gene programs, facilitating downstream regulatory network inference, functional enrichment of specific trend types, and prioritization of candidate therapeutic targets. Second, its lightweight implementation and efficient parallel computation allow it to handle large-scale datasets on standard workstations, lowering the computational and experimental barriers to adoption. Third, by providing clear trend labels and strength scores, scTrends significantly enhances the interpretability and reproducibility of results, addressing the increasing demand for transparency in single-cell genomics.

Despite these advantages, scTrends has several limitations. First, like other pseudotime-based methods, its performance is ultimately dependent on the quality and biological relevance of the input pseudotime ordering; inaccurate or overly simplified trajectories may propagate errors into trend classification. Second, although the current equal-width binning strategy is reproducible and computationally efficient, it may lose resolution in regions with highly uneven cell densities. Future versions could incorporate adaptive or density-aware binning. Third, while rule-based classification is biologically motivated, it still involves threshold selection; although scPS provides robust data-adaptive defaults, users should carefully validate results through sensitivity analysis when studying high-noise or low-cell-number trajectories. Fourth, due to the sparse nature of single-cell sequencing and the binning process, *scTrends* may struggle to identify genes with stable expression patterns, such as housekeeping genes. Even these genes can show variability, and small fluctuations beyond adaptive thresholds may occur, especially after binning and GAM smoothing. We acknowledge this limitation and will explore potential solutions in future versions of *scTrends*. Finally, while we have demonstrated its applicability in linear and partially branching scenarios, a systematic evaluation of highly complex, multi-branch trajectories (e.g., whole organ maps) remains a future direction for development.

In summary, scTrends is a robust, user-friendly, and highly interpretable tool for pseudotime-based gene expression trend analysis, filling an important gap. By linking cell ordering to actionable gene-level dynamics, it complements existing pseudotime inference methods and provides a more comprehensive, reproducible analytical approach for studying transcriptional programs in development, aging, and disease.

## Availability and requirements

Project name: scTrends (a framework for automatic classification of temporal gene expression dynamics in single-cell pseudotime analysis)

Project home page: https://github.com/746443qjb/scTrends

Operating system(s): Windows, MacOS and Linux.

Programming language: R

Other requirements:R ≥ 4.1.0 License: MIT License

Any restriction to use by non-academics: none

## List of abbreviations

ARI: Adjusted Rand Index
CV: Coefficient of Variation
GAM: Generalized Additive Model
GCV: Generalized Cross-Validation
NMI: Normalized Mutual Information
PBMC: Peripheral Blood Mononuclear Cell
RAM: Random Access Memory
scRNA-seq: Single-cell RNA Sequencing
TCM: Central Memory T Cell

## Declarations

### Ethics approval and consent to participate

Not applicable

### Consent for publication

Not Applicable

### Availability of data and materials

The scTrends package is publicly available on GitHub at https://github.com/746443qjb/scTrends, where detailed documentation and usage tutorials are also provided. The single-cell dataset used in this study is publicly available. It has been deposited in the Zenodo database under accession number 19511482. The single-cell datasets used in this study are publicly available. The human PBMC dataset can be downloaded from 10x Genomics’ Data Portal (https://seurat.nygenome.org/src/contrib/pbmcMultiome.SeuratData_0.1.4.tar.gz), the the human brain dataset is available from the CellxGene database (https://datasets.cellxgene.cziscience.com/090a448d-1dcc-4e4b-861a-6bed3867b6e0.h5ad), and the mouse pancreas dataset is available as a built-in example dataset in *scVelo*. In addition, all scripts used for data analysis and figure generation in this study are available on GitHub at https://github.com/746443qjb/scTrends_test.

### Competing interests

The authors declare no competing interests.

### Funding

This study was supported by the and the National Natural Science Foundation of China (No. 82370717), and Key Program of the Natural Science Foundation of Zhejiang Province (LZ23H050001).

### Author contributions

Conceptualization: Jianbo Qing. Methodology: Jianbo Qing. Investigation: Jianbo Qing. Validation: Jianbo Qing and Jiaying Hu. Visualization: Jianbo Qing. Formal analysis: Jianbo Qing. Funding acquisition: Junnan Wu. Project administration: Junnan Wu. Supervision: Junnan Wu. Writing—original draft: Jianbo Qing. Writing–review & editing: Xiao Wang.

## Acknowledgement

We sincerely appreciate the help provided to us during the development of *scTrends*.

## Notes

https://zenodo.org/records/19511482

### Competing Interest Statement

The authors have declared no competing interest.

