## Supplementary material for "scTrends: automated classification and strength quantification of gene expression trends along pseudotime in single-cell RNA-seq": Figure S1-S3

### Supplementary Figures

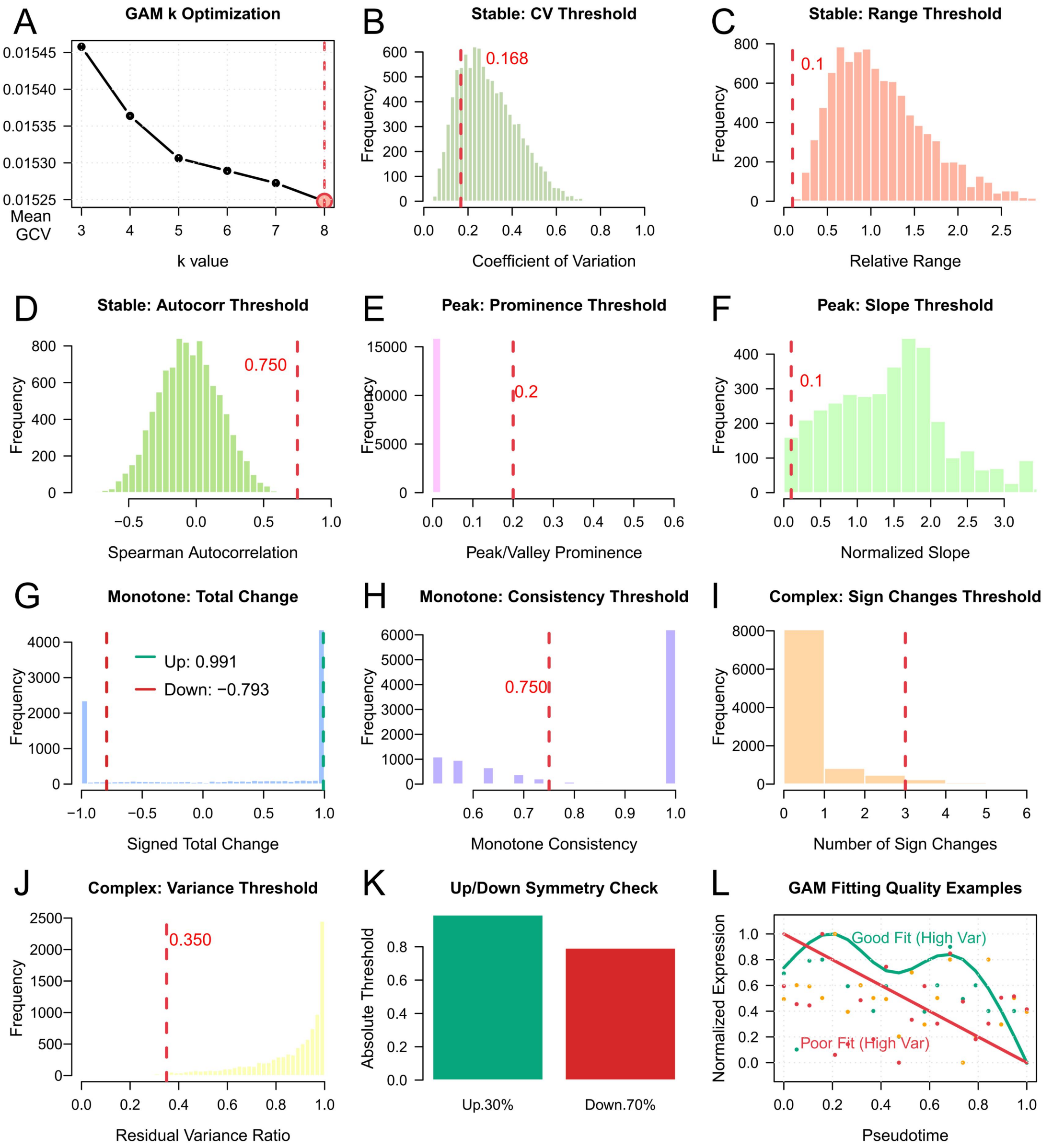


**Figure S1. Parameter selection for trends classification of PBMC dataset.** (A) Selection of the GAM spline basis dimension ($k$) based on the mean GCV score across sampled genes; the optimal $k$ is indicated by the red dashed line. (B) Distribution of the CV across sampled genes, with the selected stability threshold marked. (C) Distribution of relative expression range used for stable-gene detection. (D) Distribution of lag-1 Spearman autocorrelation coefficients, illustrating the threshold for temporal coherence in stable genes. (E) Distribution of peak/valley prominence values used to determine biologically meaningful extrema. (F) Distribution of normalized slope magnitudes around extrema, defining the minimum slope required for peak/valley classification. (G) Distribution of signed total expression change along pseudotime, with thresholds for monotone increasing (Up) and decreasing (Down) trends indicated. (H) Distribution of monotone consistency

scores, defined as the proportion of first differences sharing the same sign. (I) Distribution of the number of sign changes in first differences, used to identify complex expression patterns. (J) Distribution of residual variance ratios, quantifying unexplained variation after GAM fitting and defining the complexity threshold. (K) Symmetry check of “Up” and “Down” total-change thresholds to ensure balanced detection of increasing and decreasing trends. (L) Representative examples illustrating GAM fitting quality along pseudotime. Colored points denote normalized binned expression values for individual genes, while solid curves show the corresponding GAM fits. Different colors (green, orange, and red) represent genes with low, intermediate, and high residual variance ratios, respectively, corresponding to good, moderate, and poor model fits.

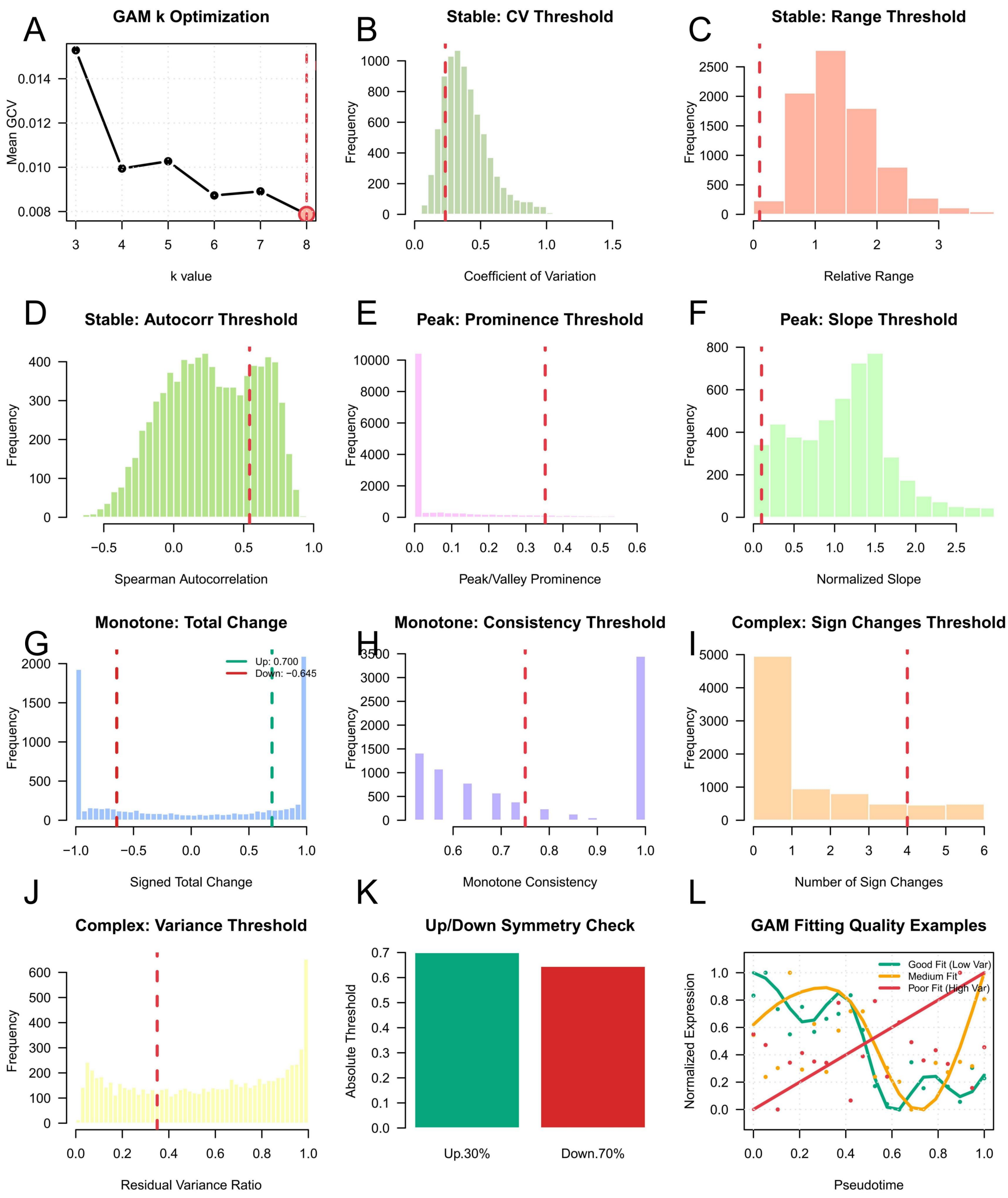


**Figure S2. Parameter selection for trends classification of oligodendrocytes dataset.** (A) Selection of the GAM spline basis dimension (*k*) based on the mean GCV score across sampled genes; the optimal *k* is indicated by the red dashed line. (B) Distribution of the CV across sampled genes, with the selected stability threshold marked. (C) Distribution of relative expression range used for stable-gene detection. (D) Distribution of lag-1 Spearman autocorrelation coefficients, illustrating the threshold for temporal coherence in stable genes. (E) Distribution of peak/valley prominence values used to determine biologically meaningful extrema. (F) Distribution of normalized slope magnitudes around extrema, defining the minimum slope required for peak/valley classification. (G) Distribution of signed total expression change along pseudotime, with thresholds for monotone increasing (Up) and decreasing (Down) trends indicated. (H) Distribution of monotone consistency

scores, defined as the proportion of first differences sharing the same sign. (I) Distribution of the number of sign changes in first differences, used to identify complex expression patterns. (J) Distribution of residual variance ratios, quantifying unexplained variation after GAM fitting and defining the complexity threshold. (K) Symmetry check of “Up” and “Down” total-change thresholds to ensure balanced detection of increasing and decreasing trends. (L) Representative examples illustrating GAM fitting quality along pseudotime. Colored points denote normalized binned expression values for individual genes, while solid curves show the corresponding GAM fits. Different colors (green, orange, and red) represent genes with low, intermediate, and high residual variance ratios, respectively, corresponding to good, moderate, and poor model fits.

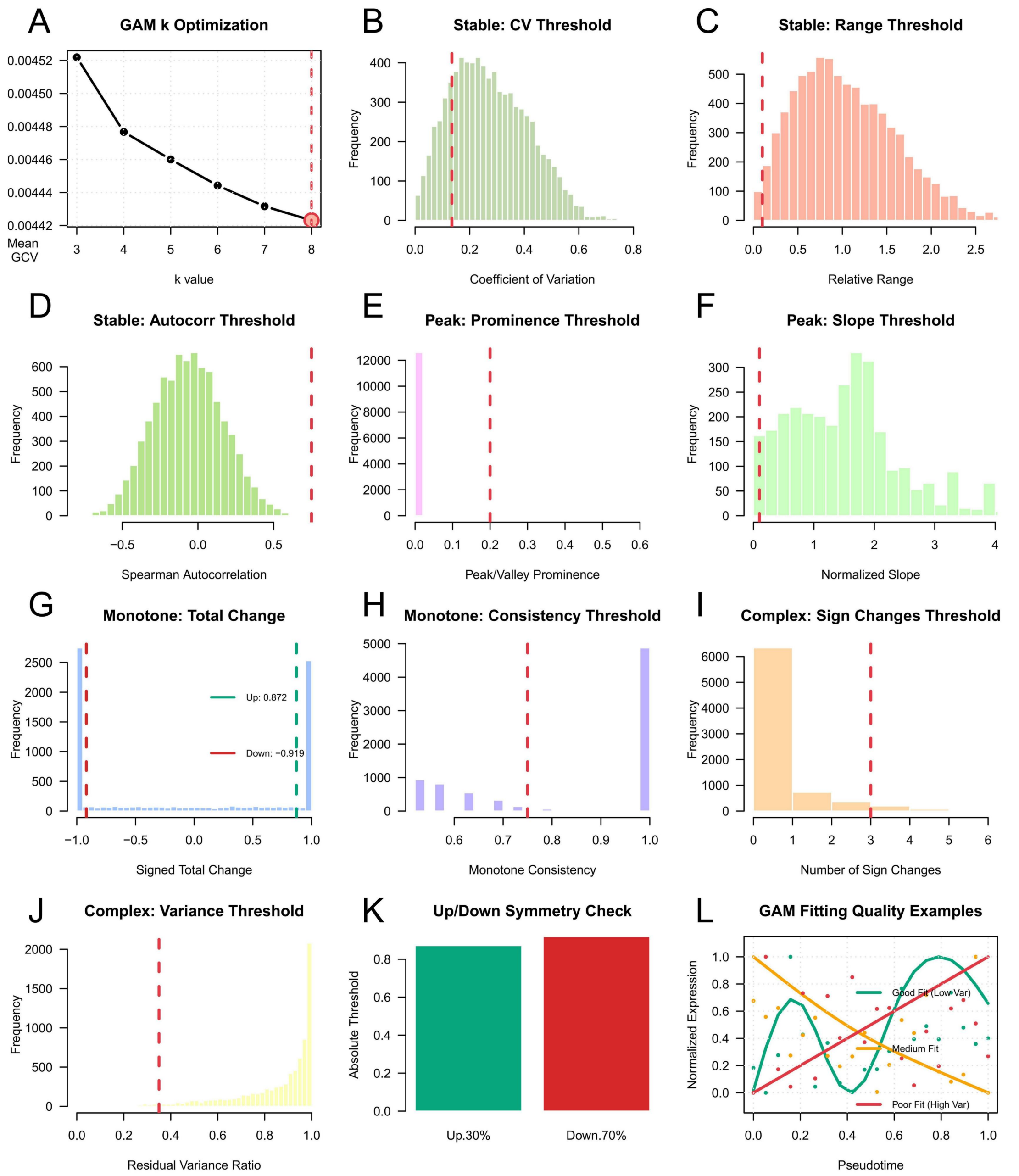


**Figure S3. Parameter selection for trends classification of pancreatic cells dataset.** (A) Selection of the GAM spline basis dimension (*k*) based on the mean GCV score across sampled genes; the optimal *k* is indicated by the red dashed line. (B) Distribution of the CV across sampled genes, with the selected stability threshold marked. (C) Distribution of relative expression range used for stable-gene detection. (D) Distribution of lag-1 Spearman autocorrelation coefficients, illustrating the threshold for temporal coherence in stable genes. (E) Distribution of peak/valley prominence values used to determine biologically meaningful extrema. (F) Distribution of normalized slope magnitudes around extrema, defining the minimum slope required for peak/valley classification. (G) Distribution of signed total expression change along pseudotime, with thresholds for monotone increasing (Up) and decreasing (Down) trends indicated. (H) Distribution of monotone consistency

scores, defined as the proportion of first differences sharing the same sign. (I) Distribution of the number of sign changes in first differences, used to identify complex expression patterns. (J) Distribution of residual variance ratios, quantifying unexplained variation after GAM fitting and defining the complexity threshold. (K) Symmetry check of “Up” and “Down” total-change thresholds to ensure balanced detection of increasing and decreasing trends. (L) Representative examples illustrating GAM fitting quality along pseudotime. Colored points denote normalized binned expression values for individual genes, while solid curves show the corresponding GAM fits. Different colors (green, orange, and red) represent genes with low, intermediate, and high residual variance ratios, respectively, corresponding to good, moderate, and poor model fits.
